# Urothelial-lineage master transcription factor hub proteomics shows mechanisms impeding urothelial cancer cell differentiation

**DOI:** 10.64898/2026.08.12.744501

**Authors:** Caroline Schuerger, Sudipta Biswas, Kwok Peng Ng, Lisa Cardone, Xiaorong Gu, Shinjini Ganguly, Rita Tohme, Arda Durmaz, Maximilian Stich, Daniel J. Lindner, Babal Jha, Omar Y. Mian, Yogen Saunthararajah

## Abstract

Urothelial cancer (UC) cells of the luminal subtype exhibit partial, incomplete differentiation towards umbrella cells that line bladder lumen, seen by morphology and gene expression. Differentiation is stalled even though the cells express master transcription factors (MTFs) that drive luminal urothelial differentiation, e.g., FOXA1 and CEBPB, at levels seen in normal differentiated urothelium. We therefore analyzed the FOXA1/CEBPB MTF hub by mass spectrometry. SWI/SNF coactivator complex (CoA) components, e.g., SMARCA4, ARID1A, that read the epigenetic activation mark histone 3 lysine 27 acetylation (H3K27ac) and use ATP-hydrolysis to ‘open’ chromatin, were the most abundant proteins pulled-down with FOXA1/CEBPB. However, genes for these and other CoA, e.g., CREBBP, EP300 that write H3K27ac, were mutated/deleted in >95% of UCs in clinical series. Also contained in the hub were corepressors (CoR) that erase H3K27ac and ‘close’ chromatin, e.g., HDAC1, CHD4 – genes for these CoR were recurrently gained in UCs. Chromatin analyses showed H3K27ac-centered remodeling was needed to activate umbrella but not constitutively accessible cell growth/division/housekeeping genes. Restoring ARID1A into *ARID1A*-mutated UC cells using lentiviral transduction, or inhibiting CoR with siRNA or small molecules, activated umbrella genes and terminated replications. In summary, UC-genesis selects for loss- and gain-of-function of CoA and CoR respectively in the urothelial-lineage MTF hub; small molecule CoR-inhibitors are candidate remedies to renew maturation towards terminal differentiated-fates.

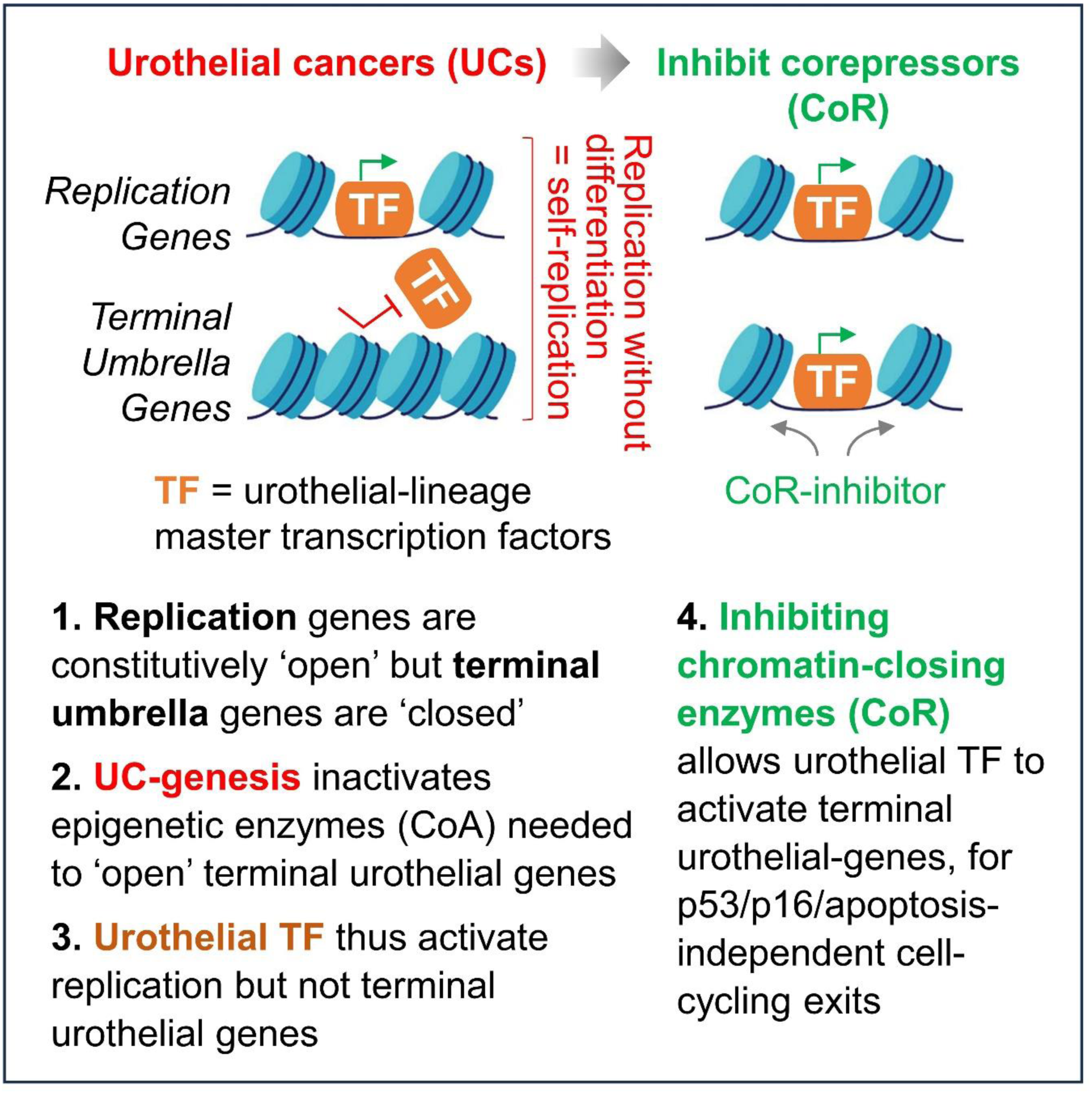

## INTRODUCTION

Most bladder cancers (∼90%) are urothelial carcinomas/cancers (UC). Gene expression signatures signifying differentiation through the stratified epithelial layers of the urinary system have been used to classify UCs as basal or luminal subtypes, with luminal subtypes being the most differentiation-advanced towards superficial umbrella cells that line the bladder lumen^1^. The greater the suppression of terminal umbrella cell genes, e.g., uroplakins, the more aggressive the UC^1–3^. There is therefore incentive to investigate mechanisms for abnormal or impeded urothelial differentiation in ways that might point to remedies.

Urothelial lineage-differentiation is driven by master transcription factors, e.g., forkhead box A1 (FOXA1), shown by: (i) FOXA1 is expressed from early through late stages of urothelial ontogeny, including high level expression in UCs^4–12^; (ii) FOXA1 directs urothelial gene expression in UC cells, seen by altered patterns of urothelial gene expression upon its introduction or deletion^11,13–15^; (iii) FOXA1 cooperates with other master transcription factors, e.g., CCAAT enhancer binding protein beta (CEBPB) which is also highly expressed in normal urothelium and UCs, to activate umbrella cell genes, e.g., uroplakins^4,11,14,16–25^; (iv) FOXA1 knock-out deranges normal bladder tissue differentiation and development^26^ (reviewed in^27^); and (iv) FOXA1 introduction can convert non-urothelial cells, e.g., fibroblasts, into urothelial-lineage cells via such cooperation (lineage-conversion)^6,9,28^. In sum, UCs are stalled in the urothelial-differentiation process even though they highly express FOXA1, CEBPB, and other master transcription factor drivers of urothelial-lineage differentiation.

FOXA1 has ‘pioneer’ functions: it can bind DNA contained in chromatin inaccessible to the basal transcription factor machinery (‘closed’ chromatin) and remodel this chromatin to create access for gene transcription^22^ (‘open’ chromatin). Chromatin remodeling for gene activation is executed by multiprotein coactivator complexes (CoAs) opposed by multiprotein corepressor (CoR) complexes that execute the opposite functions - CoAs and CoRs are recruited into master transcription factor hubs in a dynamic balance or interchange (reviewed in^29^). Combinatorial codes of master transcription factors, e.g., FOXA1 with CEBP and/or gata binding protein (GATA) family members, resolve this balance towards CoAs (CoR→CoA exchange) and thus activation of specific target genes^11,14,16,17,20,22,29^. To investigate for potential disruptions to these processes as causes of abnormal urothelial-lineage differentiation in UCs^1–3^, we immunoprecipitated endogenous FOXA1 and another lineage master transcription factor, CEBPB, from UC cells, and used liquid chromatography tandem mass spectrometry (LCMS/MS) to map recruited CoA and CoR. The results of these analyses pointed to potential mechanisms for disrupted lineage-differentiation in UCs, as well as candidate remedies to restore differentiation processes towards terminal, non-dividing fates.

## METHODS

### Cell culture

Human UC cell lines UC-6 and UC-3 were obtained from Millipore Sigma (ECACC) and cultured in Dulbecco’s Modified Eagle medium with 10% fetal bovine serum, 100U/mL penicillin and 100µg/mL streptomycin and cultured at 37°C in 5% CO2. Both cell lines were authenticated (Labcorp Cell Line Authentication, Burlington, NC) and periodically tested using the MycoAlert® Mycoplasma Detection kit (Lonza, Cat#LT07-218) to confirm they were free of mycoplasma contamination.

### Immunoprecipitation/Mass Spectrometry

Cell fractionation and nuclear protein extraction: Approximately 100 million cells per condition were used in preparation. After removal of the medium, cells were transferred to 15mL tubes and washed twice with 10mL ice-cold 1X PBS that contained protease inhibitors (Sigma-Aldrich, A8340). A total of 50µL of 10% NP-40 was added to cell suspensions to break the cell membrane. After 5-minute incubation on ice, cell suspensions were centrifuged at 344*g* for 10 minutes. The supernatant was transferred to clean 1.5mL Eppendorf tubes and labeled as the cytoplasmic fractions. Nuclear pellets were washed twice with ice-cold 1X PBS, and resuspended in 100µL of 50mM Tris-HCl, pH 8.0, 1mM MgCl2, 10mM PMSF, protease inhibitors, and Benzonase (Sigma-Aldrich D5915, 250 units). The nuclear suspensions were incubated on ice for 90 minutes with vigorous vortex every 10 minutes. At the end of incubation, 500µL protein extraction buffer (PBS + protease inhibitors) plus 500mM NaCl and 2% NP-40. After 15 minutes of incubation on ice, the mixture was centrifuged at 12396*g* for 5 minutes. The same extraction process was repeated two more times. The supernatant containing nuclear proteins was combined and transferred with clean tubes, and protein concentration was determined by BCA assay.

### Covalent binding of antibody to protein G beads

Control mouse IgG (SCBT, sc-2025) was covalently coupled to Sepharose-protein A/G beads (SCBT, sc-2003) using dimethylpimelimidate (Sigma-Aldrich, D8388). Briefly, 200µL of Sepharose-protein A/G beads was washed with 1X PBS twice, incubated with 200µL of antibody (20µg) solution (1X PBS) for 1 hour at room temperature. Antibody-bound Sepharose-protein A/G beads were then incubated with 1% bovine serum albumin in 1X PBS for 1 hour at room temperature to block nonspecific binding sites. After 3 washes with 1X PBS, 25mg of dimethylpimelimidate in 1mL of 200mM triethanolamine was added, and the coupling reaction proceeded at room temperature for 20 minutes. The reaction was repeated 2 more times with fresh addition of dimethylpimelimidate and quenched with 50mM ethanolamine. The reacted Sepharose-protein A/G beads were washed extensively with 1X PBS before immunoprecipitation. Additionally, 100 µL of HNF-3α (FOXA1) or CEBPB-conjugated Sepharose-protein A/G beads (SCBT, sc-514695 and SCBT, sc-7962) was washed with 1X PBS twice, and then incubated with 1% bovine serum albumin in 1X PBS for 1 hour at room temperature to block nonspecific binding sites. The conjugated Sepharose-protein A/G beads were then washed 3x with 1X PBS before immunoprecipitation.

### 1D SDS-polyacrylamide gel electrophoresis and Western blot analysis

Protein extracts together with molecular weight markers were subjected to 1D SDS-polyacrylamide gel electrophoresis on precast 4-12% NuPAGE gels (Invitrogen, NP0335BOX). After electrophoresis per the manufacturer’s instructions (Invitrogen), proteins were transferred to polyvinylidene difluoride membranes (Millipore) at 38 constant voltage for 1 hour using Invitrogen’s XCell II Blot module. Secondary antibodies, anti–rabbit (GE Healthcare, NA934) and anti–mouse (GE Healthcare, NXA931), were used at 1:5000 and 1:1000 dilutions, respectively. Antibodies used for detection: DNMT1 (Cell Signaling #5032), ARID1A (Sigma HPA005456 Lot 000007185).

### Protein identification by LCMS/MS

Immunoprecipitation products were subjected to SDS-polyacrylamide gel electrophoresis and stained with colloidal Coomassie Blue (Gel Code Blue, Pierce Chemical). Gel slices were excised from the top to the bottom of the lane. Proteins were reduced with 10mM dithiothreitol (Sigma-Aldrich, D0632), alkylated with 55mM iodoacetamide (Sigma-Aldrich, I1149), and digested *in situ* with trypsin. Peptides were extracted from gel pieces 3 times using 60% acetonitrile and 5% formic acid/water. The dried tryptic peptide mixture was redissolved in 20 µL of 1% formic acid for mass spectrometric analysis. Tryptic peptide mixtures were analyzed by online LC-coupled tandem mass spectrometry (LCMS/MS) on an Orbitrap mass spectrometer (Thermo Fisher Scientific).

### Database search and data validation

Mascot Daemon software (version 2.3.2, Matric Science, London UK) was used to perform database searches, using the Extract_msn.exe macro provided with Xcalibur (version 2.0 SR2, Thermo Fisher Scientific) to generate peaklists. The following parameters were set for creation of the peaklists: parent ions in the mass range 400-4500, no grouping of MS/MS scans, and a threshold at 100-. A peaklist was created for each analyzed fraction (i.e. gel slice), and individual Mascot (version 2.3.01) searches were performed for each fraction. The data was searched against *Homo sapiens* entries in Uniprot protein database (Sept 2017 release, 20,237 reviewed sequences). Carbamidomethylation of cysteines was set as a fixed modification, and oxidation of methionine was set as a variable modification. Specificity of trypsin digestion was set for cleavage after Lys or Arg, and two missed trypsin cleavage sites were allowed. The mass tolerances in MS and MS/MS were set to 10ppm and 0.6Da respectively, and the instrument setting was specified as “ESI-Trap”. To calculate the false discovery rate (FDR), the search was performed using the “decoy” option in Mascot. The spectral FDR and protein FDR are 0.42±0.09% and 4.36±1.32% respectively. A minimum Mascot ion score of 25 and peptide rank 1 was used for automatically accepting all peptide MS/MS spectra.

### Label free relative protein quantitation (LFQ)

Relative protein quantification was performed using spectral count-based LFQ. For each biological sample, data from the individual gel slices were combined. Statistical analysis was performed on all proteins identified with average spectral counts of ≥2. The spectral count data was normalized by total spectral counts of the bait protein (FOXA1 or CEBPB) in each sample to adjust for differences in overall protein levels among samples. The proteomic data has been uploaded into ProteomeXchange (accession number pending).

### Protein network analysis

Protein networks were constructed using Cytoscape 3.4. Briefly, protein-protein interactions networks were predicted using STRING with high confidence (0.70). Proteins and their normalized relative quantification value were loaded to the network as protein nodes. For individual protein interactome, the size of each protein node was formatted to be continuous mapping to the normalized relative quantification value. For comparative proteomic analysis between experiments, each protein node was presented the average of triplicate experiments.

### Lentivirus production and transduction of UC-6 cell line

Lentiviral plasmids pLenti-puro (Addgene Plasmid #39481) and pLenti-puro-ARID1A (Addgene Plasmid #39478) were packaged in HEK293T cells using second generation packaging plasmids psPAX2 and pMD2G. To increase transduction efficiency, the viral supernatant containing lentiviral particles were collected, filtered with a 0.45 μm nitrocellulose membrane and concentrated using an Amicon ultra 4. Centrifugal filter units (Millipore UFC 801024). The cells were transduced with viral particles re-suspended in 8 μg/mL polybrene for 48hrs. Post transduction the cell lines were selected with 5 μg/mL puromycin respectively 2 days, and maintained in 2 μg/mL in media with Fetal Bovine Serum-TET tested (R&D Systems S10350). Expression of transduced genes were induced by the addition of 100 µg /ml doxycycline for 48hrs.

### RNA-Seq, data processing and identification of differentially expressed genes

RNA-extraction and library preparation from the cell pellet, sequencing and data processing was performed by an external vendor, Novogene. Sequence reads were trimmed to remove possible adapter sequences and nucleotides with poor quality using Trimmomatic v.0.36. The trimmed reads were mapped to the Homo sapiens GRCh38 reference genome available on ENSEMBL using the STAR aligner v.2.5.2b. The STAR aligner is a splice aligner that detects splice junctions and incorporates them to help align the entire read sequences. BAM files were generated as a result of this step.

Unique gene hit counts were calculated by using feature Counts from the Subread package v.1.5.2. The hit counts were summarized and reported using the gene_id feature in the annotation file. Only unique reads that fell within exon regions were counted. Since strand-specific library preparation was performed, the reads were strand-specifically counted.

After extraction of gene hit counts, the gene hit counts tables, provided as Table S5, were used for downstream differential expression analysis. Using DESeq2, a comparison of gene expression between empty vector and ARID1A transduced UC-6 cells was performed. The Wald test was used to generate p-values and log2 fold changes. Genes with an adjusted p-value < 0.05 and absolute log2 fold change > 1 were called as differentially expressed genes, listed in Table S6. The raw and processed RNA-seq data are available at GEO database Accession no: GSE331331.

### siRNA knockdown of gene expression

Cells were transfected with 25 nM of either DNMT1 (Cat # 4390825, ID: s4215, ThermoFisher Scientific) or SMARCA5 (Cat # AM16704, ID: 139205, ThermoFisher Scientific) siRNA or control siRNA (Cat # 4390844, ThermoFisher Scientific) using lipofectamine RNAiMAX (Cat# 13778075, ThermoFisher Scientific) per manufacturer’s protocol. After 72 hours, knock-down was evaluated by Western blot, together with cell counts by automated cell counter, and GATA6 and UPK2 gene expression measurements by QRT-PCR.

### RNA isolation, reverse transcription (RT) and real-time PCR

Total RNA was isolated from cells using the Roche High Pure RNA Isolation Kit (#11828665001) or using RNA isolation kit (Qiagen) according to the manufacturer’s instructions. After isolation, RNA was quantified using the Nanodrop, and 1µg of RNA was used for reverse transcription reactions. The cDNA was synthesized using the iScript cDNA synthesis kit (BioRad Cat#1708891). Real time polymer chain reaction experiments were used to quantify gene expression using the ABI PRISM 7500 Fast Sequence Detection System and Fast SYBR Green Master Mix (Applied Biosystems, Cat#438610) according to the manufacturer’s instructions. Primers for all genes analyzed were purchased from Integrated DNA Technologies. The relative gene expression was analyzed following the Livak-Schmittgen method.

Primer sequences were: *ARID1A F* CTGCGTCTGTGTGTCCAATA; *ARID1A R* CGAGATGTTGGCGAGTGTAA. *B ACTIN F* ACCTTCTACAATGAGCTGCG; *B ACTIN R* CTGGATAGCAACGTACATGG. *GAPDH F* CTTTGTCAAGCTCATTTCCTGG; *GAPDH R* CTTCCTCTTGTGCTCTTGC. *UPK2 F* AATCCATTGGGCTGGGTATG; *UPK2 R* TGCCAGGGCAATGATGAA. *GATA6 F* CCAGGAAACGAAAACCTAAGAAC; *GATA6 R* TGAGGCTGTAGGTTGTGTTG

### Flow cytometry analysis

Cells were harvested after treatment and forward scatter and side scatter were acquired by flow cytometry using BD FACS Versa (BD Biosciences) and data was analyzed by FlowJo V10.

### Apoptosis assay

Cells were treated with specific drugs decitabine or CCF101 or positive control camptothecin for 24 hours. cell apoptosis was detected by caspase activity using Caspase-Glo® 3/7 Assay System from Promega (cat # G8091) following manufacturer’s instruction.

### Giemsa staining of cells

Cytospins of cells were fixed for 2 minutes in methanol/ethanol, air-dried, and stained for 20 minutes with filtered modified solution of Giemsa stain (Sigma Aldrich, Cat # 48900, St Louis, MO). Images were captured at 400X using Leica Upright Microscope-Orion.

### Bioinformatic and Statistical analysis

Urothelial genes normally most upregulated in the umbrella cells that face urine in the bladder lumen were identified by others using immunohistochemistry and single cell RNA sequencing of normal human urothelium^30,31^). Replication genes were target genes of the master transcription factor regulator of cell growth and division, identified by others using chromatin immunoprecipitation methods^32^. For comparison of gene expression between normal bladder and UC, gene expression data was downloaded from TCGA. Gene-level transcription estimates by RNA-Sequencing were analyzed as log2(x+1) transformed RSEM normalized counts. Public ChIP-seq data (FastQ files) from Encode for H3K27ac in ESC (GSM466732) were processed by UseGalaxy tools, heatmaps and signal normalization and quantification was using EASEQ tools^33^, aligned ChIP-Seq reads were normalized to reads per million per 1kbp. Mann-Whitney and Student’s t-tests were 2-sided and performed at least the 0.05 significance level or at even more stringent levels of significance as stated. SAS statistical software (SAS Institute Inc., Cary, NC) or PRISM software (GraphPad, San Diego, CA) was used to perform statistical analyses.

## RESULTS

### CoAs recruited by FOXA1/CEBPB are recurrently mutated/deleted while CoRs are amplified

Endogenous FOXA1 and CEBPB were immunoprecipitated from UC cells (UC-6) each in triplicate experiments, using immunoprecipitations with IgG isotype controls in quadruplicate as controls, and recruited proteins were identified by liquid chromatography tandem mass spectrometry (IP-LCMS/MS). Comparing total spectral count data for FOXA1 and CEBPB versus IgG isotype immunoprecipitations identified components of the SWI/SNF BAF CoA complex, e.g., AT-rich interaction domain 1A (ARID1A), as the proteins most abundantly pulled-down with FOXA1 or CEBPB (Benjamani-Hochberg adjusted p-values <0.05, **Table S1**, **S2**) (**Figure 1A**). Key members of the SWI/SNF PBAF complex were not detected. The SWI/SNF complex reads the histone 3 lysine 27 acetylation (H3K27ac) epigenetic activation mark and uses ATP-hydrolysis to reposition nucleosomes; relatedly, CREB binding lysine acetyltransferase (CREBBP) and EP300 lysine acetyltransferase (EP300) that write H3K27ac were also found in the FOXA1 and CEBPB interactomes (**Figure 1B**).

**Figure 1.**
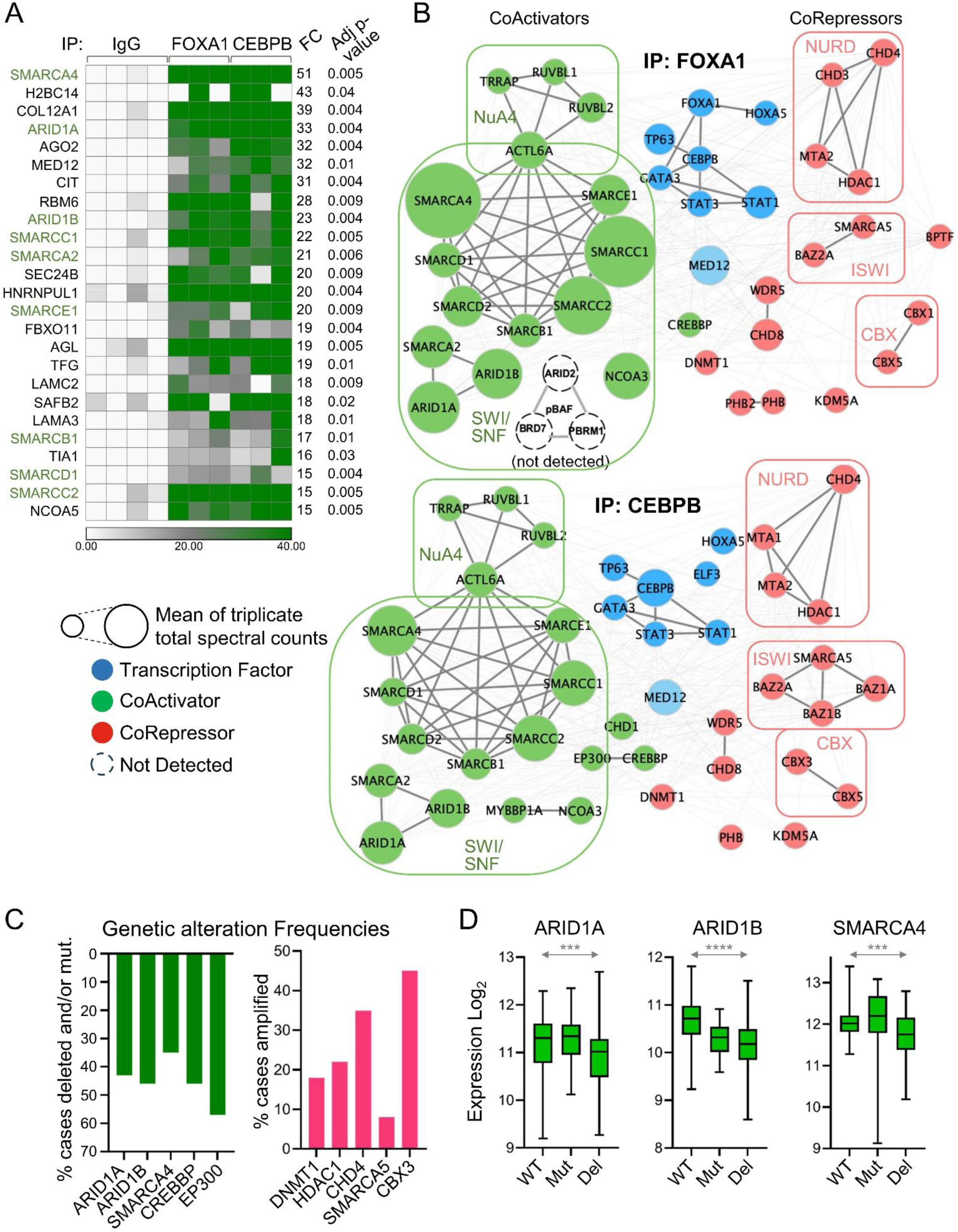
Coactivators (CoA) and corepressors (CoR) recruited by endogenous FOXA1 and CEBPB in UC cells are recurrently genetically altered. **A) Total spectral count data for FOXA1 and CEBPB versus IgG isotype immunoprecipitations (IPs) identified components of the SWI/SNF BAF CoA complex, e.g., ARID1A, as the proteins most abundantly pulled-down with FOXA1 or CEBPB** (Benjamani-Hochberg adjusted p-values <0.05, **Table S1**, **S2**). FC=fold-change IP: FOXA1/CEBPB vs IgG, Benjamani-Hochberg adjusted p-values, full data in **Table S1**, **S2**. Key members of the SWI/SNF PBAF complex were not detected. SWI/SNF members highlighted with green font. **B) Map of CoA and CoR pulled down with FOXA1 and CEBPB.** Endogenous FOXA1 or CEBPB were IPed from UC-6 cells and the interactome analyzed by LCMS/MS in triplicate; CoA (green), CoR (red), and transcription factors (dark blue) in the interactomes are shown. A minimum Mascot ion score of 25 and peptide rank 1 was used for automatically accepting all peptide MS/MS spectra. Circle size indicates abundance of protein in the interactome, average of triplicate IP-LCMS/MS experiments. Known coregulator multi-protein complexes are grouped together. Spectral counts **Table S1**, differential analyses versus IP with IgG isotype control in **Table S2**. **C) CoA recruited by FOXA1/CEBPB were recurrently mutated and/or deleted while CoR were recurrently amplified** (analyses of TCGA database^34^). Gene-level copy number estimates generated by the GISTIC2 method were thresholded to estimated values with -2 and -1 representing homozygous or single copy deletion (Del), 0 representing diploid normal copy, and 1 or 2 representing low-level or high-level copy number amplification (Amp). Mutation calling also as per TCGA. UCs n=397. **D) Copy number loss in CoA genes significantly decreased their expression**. Gene-level transcription estimates by RNA-Sequencing were analyzed as log2(x+1) transformed RSEM normalized counts. Median+/-interquartile range, Mann-Whitney test 2-sided, *p<0.05, **p<0.01, ***p<0.001, ****p<0.0001. Additional data in **Figure S1**.

The most prominent CoR pulled down with the master transcription factors were DNA methyltransferase 1 (DNMT1, the maintenance methyltransferase but also a CoR recruited into lineage master transcription factor hubs) and components of NuRD, ISWI and CBX complexes including histone deacetylase 1 (HDAC1), chromodomain helicase DNA binding protein 4 (CHD4), SNF2 related chromatin remodeling ATPase 5 (SMARCA5), and chromobox 3 (CBX3) that functionally oppose SWI/SNF and/or erase H3K27ac (**Figure 1B**).

Other master transcription factors known to have essential roles in urothelial-lineage differentiation, e.g., GATA3, also pulled-down with FOXA1 or CEBPB (**Figure 1B**).

We then analyzed the genetic status of these CoA and CoR in UCs using The Cancer Genome Atlas (TCGA) database^34^. Approximately 98% of UCs deleted and/or mutated one or more of *ARID1A*, *ARID1B*, *SMARCA4*, *CREBBP* or *EP300* (**Figure 1C, S1A**). The copy number losses in these CoA genes in UCs produced significant reductions in gene expression versus UCs without these alterations (**Figure 1D**). Conversely, genes for CoR components DNMT1, HDAC1, CHD4, SMARCA5 and CBX3 were recurrently gained or amplified, with ∼80% of cases gaining or amplifying ≥1 of these genes (**Figure 1C, S1B**). The CoR gene gains/amplifications were accompanied by significantly higher CoR gene expressions (**Figure S1C**).

### Chromatin remodeling requirements and expression of umbrella versus replication genes

Genes normally activated with urothelial-lineage differentiation into umbrella cells (‘*umbrella genes*’), and genes that mediate cell growth and division (‘*replication genes*’) which are suppressed upon umbrella cell differentiation, have been curated in the literature^30–32^ (**Figure 2A**, **Table S3**). As expected, umbrella-gene expression was decreased in UCs compared to non-cancerous bladder, while replication-gene expression was increased (analyses of TCGA data) (**Figure 2B, S2, Table S4**). Previous reports have indicated that the greater the suppression of terminal umbrella cell genes, the more aggressive the UC^1–3^, and consistent with this, lower umbrella gene expression in UCs correlated significantly with higher replication gene expression (**Figure 2C**).

**Figure 2.**
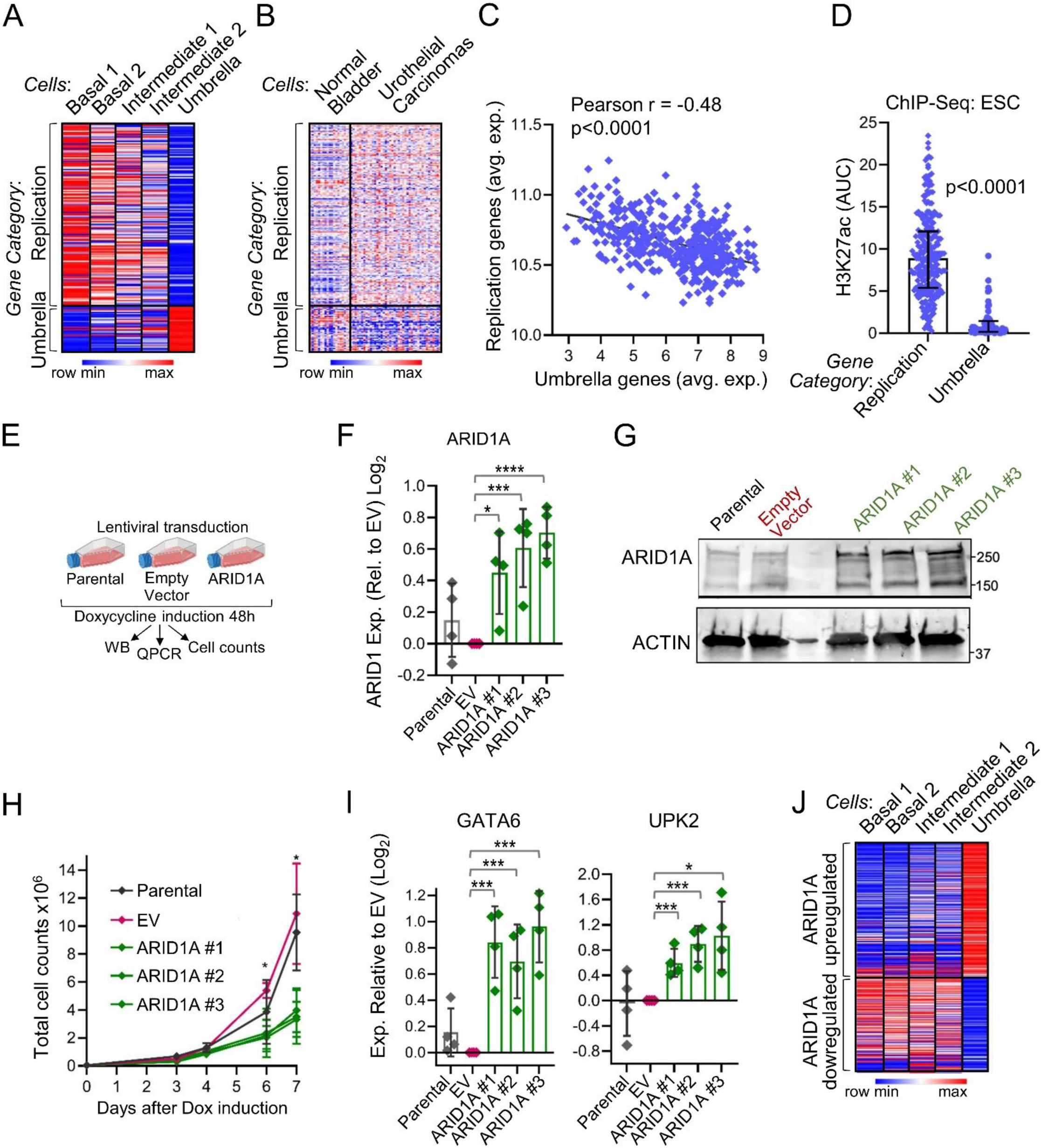
Chromatin remodeling requirements and expression of umbrella genes versus replication genes - restoring ARID1A into *ARID1A*-mutated UC cells activated umbrella genes and terminated replications. **A) Umbrella and replication gene expression in basal, intermediate, and umbrella strata of normal urothelium.** Known umbrella and replication genes (**Table S3**)^30–32^ were examined for their gene expression in a database of single cell RNA-sequencing of normal human bladder^31^. **B) Umbrella gene expression was lower, and replication gene expression higher, in primary UC tissue compared to non-cancerous bladder tissue** (TCGA RNA-sequencing, **Table S4**, non-cancer bladder n=19, primary UC n=407 truncated, full heatmap in Figure S2. **C) Less umbrella gene expression correlated with higher replication gene expression.** Primary UCs as per panel B and Figure S2. **D) H3K27ac peaks quantified at replication versus umbrella genes at the tissue baseline of embryonic stem cells** (H3K27ac is written and read by CREBBP, EP300, ARID1A, that are recruited by FOXA1 and recurrently deleted in UCs). Public ChIP-seq data, Encode GEO Database GSM466732^11^. FastQ files processed using UseGalaxy tools. Signal normalization and peak quantification by EASEQ tools^33^; p-values Mann-Whitney test, 2-sided. E) ARID1A was introduced into UC-6 cells that contain an inactivating mutation in *ARID1A*^35^. Puromycin-selectable lentiviral system containing a doxycycline-inducible ARID1A cassette versus Empty cassette. Transductions and selection of stable transductants with puromycin performed in triplicate. ARID1A expression was induced by. **F) ARID1A mRNA expression**, after 48 hours doxycycline 100µg/mL. Parenteral cells, Empty Vector transduced clone and 3 separately selected ARID1A-transduced clones. QRT-PCR quadruplicate per clone. Unpaired t-test, 2-tailed, *p<0.05, ***p<0.001, ****p<0.0001. **G) ARID1A protein.** Measured by Western blot, cells per previous panel. **H) ARID1A, but not Empty cassette, significantly decreased proliferation.** Cell counts by automated counter beginning with doxycyline addition. Unpaired t-test, 2-tailed, *p<0.05. **I) Umbrella cell marker gene GATA6 and UPK2 expression.** QRT-PCR, **c**ells as per mRNA/Western blot panels above. Unpaired t-test, 2-tailed, *p<0.05, ***p<0.001. **J) Genes significantly up- and down-regulated in ARID1A versus Empty Vector transduced** UC6 cells were in large part genes up- and down-regulated with urothelial differentiation into umbrella cells. Genes differentially expressed in ARID1A vs Empty Vector transduction identified by RNA-sequencing and DESeq2 analyses (**Tables S5**, **S6**, FDR <0.05) were mapped into a database of normal human bladder single cell RNA-sequencing to understand how their expression changes during normal urothelial differentiation^31^.

We examined if there were differential requirements for H3K27ac related remodeling at umbrella versus replication genes. At the tissue ontogeny baseline of embryonic stem cells, replication genes (like other housekeeping genes) were enriched for H3K27ac, contrasting with scarcity of H3K27ac at umbrella genes (**Figure 2D**). These results indicated a particular need for H3K27ac related chromatin remodeling to activate umbrella versus replication genes.

### Reintroducing ARID1A into ARID1A-mutated UC cells activated umbrella genes and terminated replications

To experimentally determine if CoA restoration would activate umbrella genes, we restored ARID1A into UC-6 cells that contain an inactivating mutation in *ARID1A*^35^ using a lentiviral doxycycline-inducible transduction system (**Figure 2E**). ARID1A mRNA and protein induction by 48 hours of doxycycline was confirmed in 3 separately selected ARID1A-transduced clones, and not observed in empty-vector transduced controls, measured by QRT-PCR and Western blots (**Figure 2F, G**). The ARID1A, but not Empty Vector, transduction significantly decreased proliferation of the UC cells (**Figure 2H**), accompanied by significant upregulation of umbrella cell marker genes *GATA6* and uroplakin 2 (*UPK2)* measured by QRT-PCR (**Figure 2I**). To expand the gene expression analyses, RNA-sequencing was performed on the Empty Vector and ARID1A-transduced clones (**Table S5**). Differentially expressed genes (DESeq2, FDR <0.05, **Table S6**) were annotated for links to urothelial-lineage differentiation (basal→intermediate→umbrella) by mapping into a database of normal human bladder single cell RNA-sequencing: genes significantly upregulated with ARID1A transduction were genes normally upregulated with umbrella cell differentiation, whereas >50% of the genes downregulated with ARID1A transduction were genes normally downregulated with umbrella cell differentiation (**Figure 2J**).

### Depleting the CoR DNMT1 from UC cells activated umbrella genes and decreased replications

DNMT1 is a CoR found in the FOXA1/CEBPB hub and recurrently amplified in UCs. We used siRNA to knock-down DNMT1 from UC-3 cells (**Figure 3A**). The DNMT1-depletion increased expression of the umbrella cell marker genes *UPK2* and *GATA6* measured by QRT-PCR (as also observed with ARID1A reintroduction) (**Figure 3A**), and decreased UC-3 cell proliferation (**Figure 3A**). DNMT1 can also be depleted using the small molecule decitabine, confirmed in UC-6 and UC-3 cells (**Figure 3B**). DNMT1 depletion by decitabine also activated *UPK2* and *GATA6* (**Figure 3C**). Supporting resumed urothelial-lineage maturation, cell morphology changed with increased cell size and cytoplasmic granularity, seen by light microscopy and by flow cytometry measurement of forward- and side-scatter (**Figure 3D, E**). Also as expected with lineage-maturation, cell numbers significantly decreased with decitabine versus vehicle treatment (**Figure 3F**). The decrease in cell numbers was not because of apoptosis (no apoptosis detected by caspase 3/7 assay)(**Figure 3G**).

**Figure 3.**
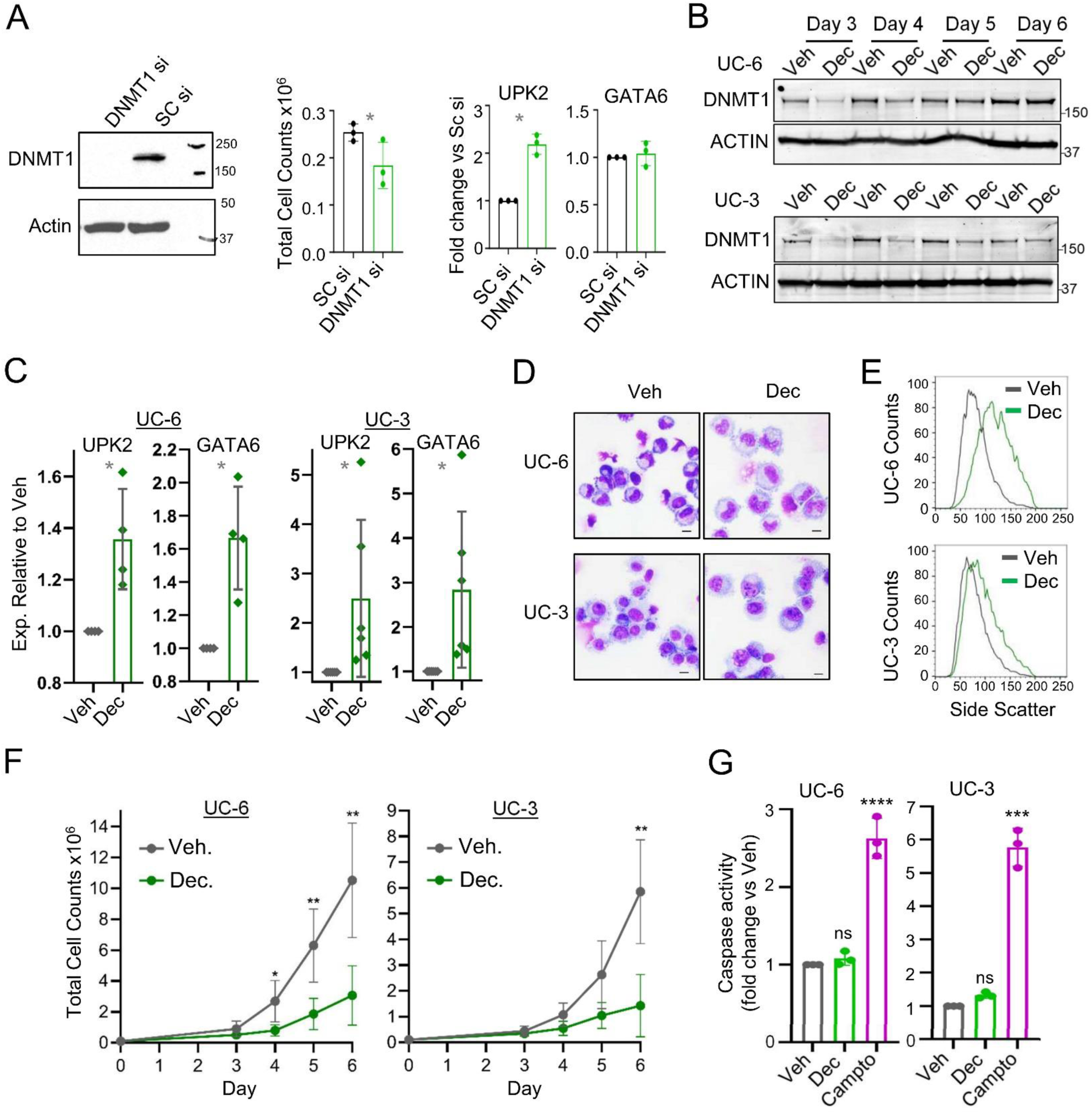
Inhibiting the CoR DNMT1 in UC cells using siRNA or decitabine (Dec) activated umbrella marker genes and decreased proliferation. **A) siRNA knockdown of DNMT1 activated umbrella genes and decreased proliferation**. UC-3 cells were transfected with 25 nM of either siDNMT1 or control siRNA using lipofectamine. After 72 hours, knock-down was evaluated by Western blot, together with cell counts by automated cell counter, and GATA6 and UPK2 gene expression measurements by QRT-PCR. Paired t-test, 2-sided, *p<0.05. **B) DNMT1-depletion by Dec treatment was confirmed by Western blot**. UC-6 and UC-3 cells were treated with Veh or Dec 0.5 µM. **C) DNMT1-targeting by Dec activated UPK2 and GATA6**. UC-6: 4 independent experiments; UC-3: 6 independent experiments. Unpaired t-test, 2-sided, *p<0.05. **D) The DNMT1-targeting increased UC nuclear and cell size and increased cytoplasmic granularity.** Giemsa-stained cytospin preparations of cells collected after 96h of treatment. Magnification 400X. Scale bar 20 μm. **E) The DNMT1-targeting increased UC cell granularity (side-scatter).** Cell granularity was accessed by flow cytometry and data shown as histograms for side scatter. **F) DNMT1-targeting by Dec decreased UC-6 and UC-3 cell proliferation**. Cell growth was compared Veh vs Dec 0.5µM treatment. Cumulative cell counts by automated counter after treatment for indicated time points. Mean±standard deviation (SD) for 4 independent experiments. Unpaired t-test, 2-sided, *p<0.05, **p<0.01. **G) Anti-proliferative effect of Dec was not via apoptosis.** Apoptosis was measured by caspase activity assay after 24 hours of treatment. Camptothecin 20 μM was used as positive control. one-way ANOVA test, ***p<0.001, ****p<0.0001, ns p>0.05.

### Inhibiting ISWI/CHD CoRs also activated umbrella genes and decreased replications

Also in the FOXA1/CEBPB hub were CoR of imitation switch (ISWI) and chromodomain helicase DNA binding (CHD) families, SMARCA5 and CHD4 respectively, that directly oppose SWI/SNF CoA - ISWI/CHD use energy from ATP hydrolysis to reposition nucleosomes in proximity to gene transcription start sites (‘close’ chromatin).

We used siRNA to knock-down SMARCA5 from UC-3 cells (**Figure 4A**). The SMARCA5-depletion increased expression of the umbrella cell marker genes *UPK2* and *GATA6* measured by QRT-PCR (as also observed with ARID1A reintroduction) (**Figure 4A**), and decreased UC-3 cell proliferation (**Figure 4A**). A small molecule tool compound to inhibit ISWI and CHD family functions is available, CCF101^36–40^. CCF101 treatment of UC-6 and UC-3 cells activated the umbrella cell signature genes *UPK2* and *GATA6* several-fold (**Figure 4B**), accompanied by morphology changes of urothelial-lineage maturation (decreased nuclear cytoplasmic ratio, increased cytoplasmic complexity) shown by Giemsa-staining and by flow cytometry (**Figure 4C, D**). The CCF101 treatment markedly decreased proliferation of the cells (**Figure 4E**). This proliferation-inhibition was not by apoptosis as measured by caspase activity after 24 hours of drug treatment (**Figure 4F**).

**Figure 4.**
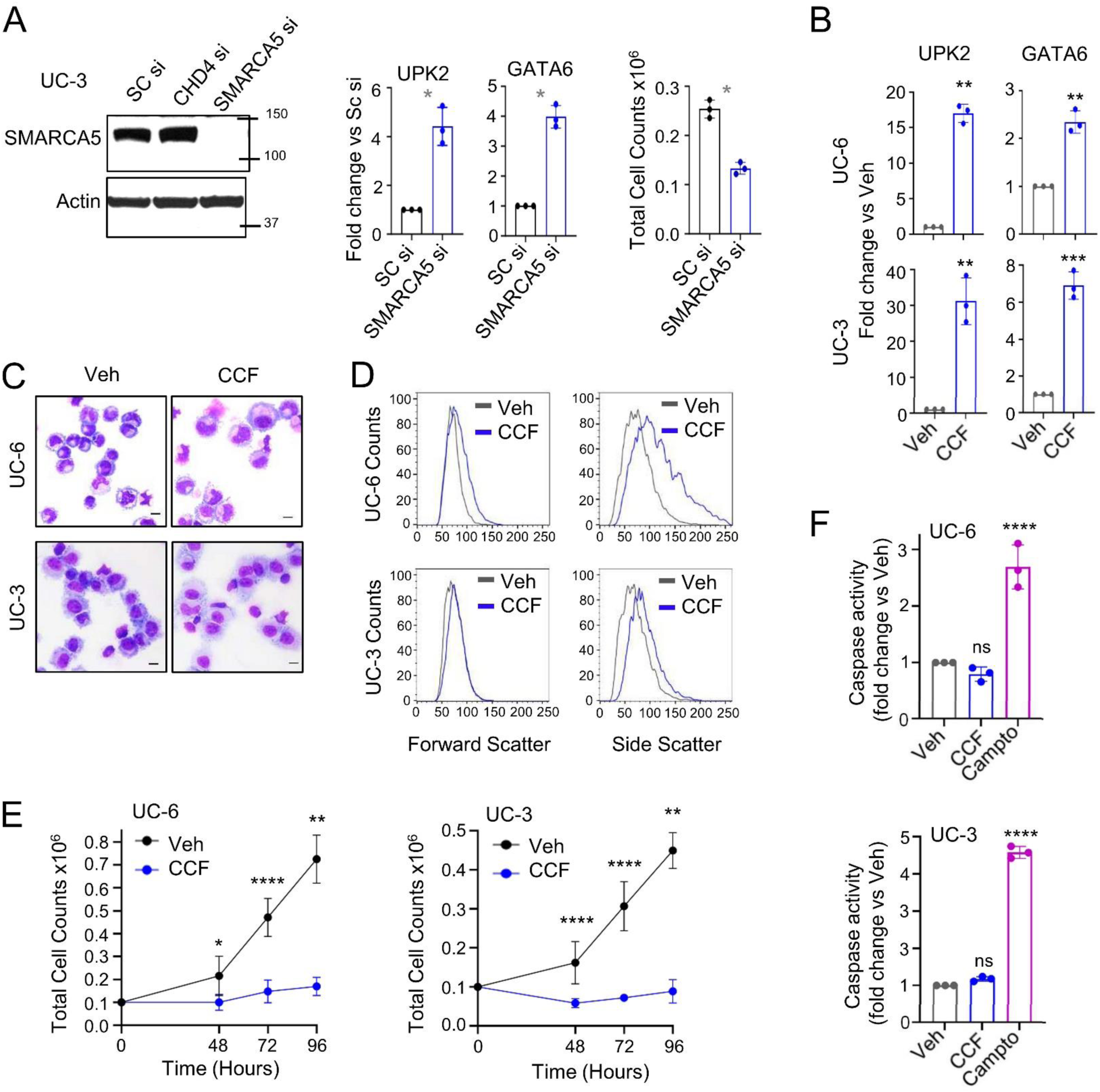
Inhibiting SMARCA5 (ISWI family CoR) with siRNA or a small molecule activated umbrella genes and decreased proliferation. **A) siRNA knockdown of SMARCA5 activated umbrella genes and decreased proliferation**. UC-3 cells were transfected with 25 nM of either siSMARCA5 or control siRNA using lipofectamine. After 72 hours, knock-down was evaluated by Western blot, together with cell counts by automated cell counter, and GATA6 and UPK2 gene expression measurements by QRT-PCR. CHD4 is a CoR in the CHD family phylogenetically related to the ISWI family. Paired t-test, 2-sided, *p<0.05. **B) A small molecule inhibitor of SMARCA5 and CHD4 activated umbrella signature genes UPK2 and GATA6**. QRT-PCR after 72 hours of treatment of UC-6 and UC-3 cells with 10 μM CCF101 (tool compound that inhibits SMARCA5 and CHD4). 3 independent experiments. Unpaired t-test, 2-sided, **p<0.01, ***p<0.001. **C) Cell morphology**. Giemsa-stained cell cytospin preparations were imaged by Leica Upright Microscope-Orion after 96h of treatment. Magnification 400X, scale bar 20 μm. **D) Cell size (forward scatter) and granularity (side-scatter).** Cell size and granularity measured by flow cytometry at 96h of treatment. **E) Cell counts with SMARCA5/CHD4-inhibitor.** UC-6 and UC-3 cells were treated with 10 μM CCF101 and cell count and viability were measured by automated cell counter from 48h to 96h. Unpaired t-test 2-sided, *p<0.05, **p<0.01, ****p<0.0001. **F) Apoptosis was not activated**. Caspase activity assay after 24 hours of treatment. Camptothecin (20 μM) was used as a positive control for apoptosis. Mean±SD of three independent experiments. Unpaired t-test, 2-sided, ****p<0.0001.

## DISCUSSION

Proteomic analyses of the FOXA1/CEBPB urothelial lineage master transcription factor hub in UC cells demonstrated major recruitment of the SWI/SNF CoA complex, e.g., SMARCA4, ARID1A, that reads the H3K27ac mark and repositions nucleosomes for gene activation. Mass spectrometry analyses of the FOXA1 interactome in MCF7 breast cancer cells by others also identified ARID1A, EP300 and CREBBP as amongst the most prominently recruited proteins^41^. These CoA are highly recurrently deleted/mutated in UCs and other cancers. Epigenetic mark analyses indicated that the chromatin remodeling functions executed by the CoA are particularly needed to activate late-lineage, e.g., umbrella genes, but not replication/housekeeping genes. Reintroduction of ARID1A into an ARID1A-mutated UC cell line activated umbrella cell genes and terminated replications. A cause-effect relationship between CoA mutations/deletions and impeded urothelial-lineage differentiation is also supported by data from others: (i) knock-out of *Arid1a* from murine urothelial precursors repressed the umbrella cell marker gene *Upk3a* and increased urothelial precursor proliferation^42^ and; (ii) in primary UC tissue, SWI/SNF CoA complex expression was lowest in least differentiated regions, quantified by immunohistochemistry^43^.

While CoA components recruited by FOXA1/CEBPB were recurrently deleted in UCs with correspondingly lower expression, specific CoRs recruited by FOXA1/CEBPB, e.g., DNMT1, CHD4, SMARCA5, CBX3, were recurrently amplified (gain-of-function) and more highly expressed. In vitro, knocking-down DNMT1 or SMARCA5 with siRNA, or their inhibition with small molecules, activated umbrella cell genes, produced morphologic changes of terminal-maturation, and decreased UC replications. Others have observed that inhibiting HDAC CoRs with small molecules also terminates UC cell replications, again with phenotype changes consistent with renewed urothelial lineage-differentiation^44,45^. Cell cycling exits by resumed lineage-maturations do not require the p53/p16-regulated apoptosis system utilized by chemo-radiation, and small molecule CoR-inhibitors can thus cytoreduce p53/p16-attenuated chemo-resistant cancer cells (reviewed in^29,46^).

A known motif in gene regulation is cooperation between master transcription factors, e.g., FOXA1 with CEBP and/or GATA family members, to produce CoR→CoA exchange and thus chromatin remodeling for gene activation^16–18,47,48^. Thus, recurrent haploinsufficiency of GATA family members, e.g., *GATA4*, which is another genetic feature of UCs, may conceivably also contribute to CoR/CoA imbalance in the urothelial-lineage master transcription factor hub^16–18,47,48^.

### Study limitations

One limitation of this study is that the proteomic analyses, and the cause-effect analyses using CoA restoration or CoR inhibition, were performed in cell lines, as is done widely for practical reasons. To mitigate this limitation, corroborating data was obtained from analyses of large databases of primary UC clinical series, and from experimental data published by others. Translation of the mechanistic, cause-effect conclusions into clinical therapy for UCs will require overcoming fundamental pharmacologic limitations of presently available CoR-inhibitors that we and others tested - DNMT1-inhibitors decitabine and azacitidine distribute preferentially into hematopoietic tissues, and HDAC-inhibitors have significant clinical on-target toxicities (HDACs have pleiotropic cell physiology roles outside of chromatin (reviewed in^29,46,49^) - and/or development of novel CoR-inhibitors, e.g., the ISWI/CHD-inhibitor tested here.

Clinical aggression and disrupted umbrella cell gene activation in UCs track together^1–3^, motivating study of mechanisms impeding urothelial differentiation. Here, we found that CoR/CoA imbalance in the urothelial lineage master transcription factor hub is one such mechanism; small molecule CoR-inhibitors are candidate remedies.

#### Authors’ Contributions

**Conception and design:** Y. Saunthararajah

**Development of methodology:** C. Schuerger, S. Biswas, X. Gu, S. Ganguly, R. Tohme, D. Lindner, B. Jha, O. Mian, Y. Saunthararajah

**Acquisition of data:** C. Schuerger, S. Biswas, L. Cardone, KP. Ng, X. Gu, S. Ganguly, R. Tohme, D. Lindner, B. Jha, O. Mian, Y. Saunthararajah

**Analysis and interpretation of data:** C. Schuerger, S. Biswas, X. Gu, A. Durmaz, M. Stich, O. Mian, Y. Saunthararajah

**Writing, review, and/or revision of the manuscript:** C. Schuerger, S. Biswas, L. Cardone, KP. Ng, X. Gu, S. Ganguly, R. Tohme, A. Durmaz, M. Stich, D. Lindner, B. Jha, O. Mian, Y. Saunthararajah

**Administrative, technical, or material support:** Y. Saunthararajah

**Study supervision:** B. Jha, O. Mian, Y. Saunthararajah

## REFERENCES

1. DeGraff DJ, Cates JM, Mauney JR, Clark PE, Matusik RJ, Adam RM. When urothelial differentiation pathways go wrong: implications for bladder cancer development and progression. Urol Oncol. 2013;31(6):802–811.

2. Humphrey PA, Moch H, Cubilla AL, Ulbright TM, Reuter VE. The 2016 WHO Classification of Tumours of the Urinary System and Male Genital Organs-Part B: Prostate and Bladder Tumours. Eur Urol. 2016;70(1):106–119.

3. Duex JE, Swain KE, Dancik GM, et al. Functional Impact of Chromatin Remodeling Gene Mutations and Predictive Signature for Therapeutic Response in Bladder Cancer. Mol Cancer Res. 2018;16(1):69–77.

4. Fishwick C, Higgins J, Percival-Alwyn L, et al. Heterarchy of transcription factors driving basal and luminal cell phenotypes in human urothelium. Cell Death Differ. 2017;24(5):809–818.

5. Bernardo GM, Keri RA. FOXA1: a transcription factor with parallel functions in development and cancer. Biosci Rep. 2012;32(2):113–130.

6. Oottamasathien S, Wang Y, Williams K, et al. Directed differentiation of embryonic stem cells into bladder tissue. Dev Biol. 2007;304(2):556–566.

7. Varley CL, Bacon EJ, Holder JC, Southgate J. FOXA1 and IRF-1 intermediary transcriptional regulators of PPARgamma-induced urothelial cytodifferentiation. Cell Death Differ. 2009;16(1):103–114.

8. Lupien M, Eeckhoute J, Meyer CA, et al. FoxA1 translates epigenetic signatures into enhancer-driven lineage-specific transcription. Cell. 2008;132(6):958–970.

9. Inoue Y, Kishida T, Kotani SI, et al. Direct conversion of fibroblasts into urothelial cells that may be recruited to regenerating mucosa of injured urinary bladder. Sci Rep. 2019;9(1):13850.

10. Reddy OL, Cates JM, Gellert LL, et al. Loss of FOXA1 Drives Sexually Dimorphic Changes in Urothelial Differentiation and Is an Independent Predictor of Poor Prognosis in Bladder Cancer. Am J Pathol. 2015;185(5):1385–1395.

11. Neyret-Kahn H, Fontugne J, Meng XY, et al. Epigenomic mapping identifies an enhancer repertoire that regulates cell identity in bladder cancer through distinct transcription factor networks. Oncogene. 2023;42(19):1524–1542.

12. Thomas JC, Oottamasathien S, Makari JH, et al. Temporal-spatial protein expression in bladder tissue derived from embryonic stem cells. J Urol. 2008;180(4 Suppl):1784–1789.

13. Kim E, Choi S, Kang B, et al. Creation of bladder assembloids mimicking tissue regeneration and cancer. Nature. 2020;588(7839):664–669.

14. Warrick JI, Walter V, Yamashita H, et al. FOXA1, GATA3 and PPARɣ Cooperate to Drive Luminal Subtype in Bladder Cancer: A Molecular Analysis of Established Human Cell Lines. Sci Rep. 2016;6:38531.

15. Iyyanki T, Zhang B, Wang Q, et al. Subtype-associated epigenomic landscape and 3D genome structure in bladder cancer. Genome Biol. 2021;22(1):105.

16. Enane FO, Shuen WH, Gu X, et al. GATA4 loss of function in liver cancer impedes precursor to hepatocyte transition. J Clin Invest. 2017;127(9):3527–3542.

17. Takaku M, Grimm SA, De Kumar B, Bennett BD, Wade PA. Cancer-specific mutation of GATA3 disrupts the transcriptional regulatory network governed by Estrogen Receptor alpha, FOXA1 and GATA3. Nucleic Acids Res. 2020;48(9):4756–4768.

18. Mauney JR, Ramachandran A, Yu RN, Daley GQ, Adam RM, Estrada CR. All-trans retinoic acid directs urothelial specification of murine embryonic stem cells via GATA4/6 signaling mechanisms. PLoS One. 2010;5(7):e11513.

19. Eriksson P, Aine M, Veerla S, Liedberg F, Sjodahl G, Hoglund M. Molecular subtypes of urothelial carcinoma are defined by specific gene regulatory systems. BMC Med Genomics. 2015;8:25.

20. Xu C, Kleinschmidt H, Yang J, et al. Systematic dissection of sequence features affecting binding specificity of a pioneer factor reveals binding synergy between FOXA1 and AP-1. Mol Cell. 2024;84(15):2838–2855 e2810.

21. Yee CH, Zheng Z, Shuman L, et al. Maintenance of the bladder cancer precursor urothelial hyperplasia requires FOXA1 and persistent expression of oncogenic HRAS. Sci Rep. 2019;9(1):270.

22. Cirillo LA, Lin FR, Cuesta I, Friedman D, Jarnik M, Zaret KS. Opening of compacted chromatin by early developmental transcription factors HNF3 (FoxA) and GATA-4. Mol Cell. 2002;9(2):279–289.

23. Nishimura Y, Sasagawa S, Ariyoshi M, et al. Systems pharmacology of adiposity reveals inhibition of EP300 as a common therapeutic mechanism of caloric restriction and resveratrol for obesity. Front Pharmacol. 2015;6:199.

24. Tomizawa M, Shinozaki F, Motoyoshi Y, Sugiyama T, Yamamoto S, Ishige N. Transcription Factors and Medium Suitable for Initiating the Differentiation of Human-Induced Pluripotent Stem Cells to the Hepatocyte Lineage. J Cell Biochem. 2016;117(9):2001–2009.

25. Adib E, Nassar AH, Abou Alaiwi S, et al. Epigenetic Atlas of Bladder Cancer Reveals Master Transcription Factors and Risk-Associated Regulatory Elements in Luminal and Basal-Squamous Molecular Subtypes. Mol Cancer Res. 2026;24(6):504–517.

26. Gredler ML, Patterson SE, Seifert AW, Cohn MJ. Foxa1 and Foxa2 orchestrate development of the urethral tube and division of the embryonic cloaca through an autoregulatory loop with Shh. Dev Biol. 2020;465(1):23–30.

27. Ramal M, Corral S, Kalisz M, Lapi E, Real FX. The urothelial gene regulatory network: understanding biology to improve bladder cancer management. Oncogene. 2024;43(1):1–21.

28. Osborn SL, Thangappan R, Luria A, Lee JH, Nolta J, Kurzrock EA. Induction of human embryonic and induced pluripotent stem cells into urothelium. Stem Cells Transl Med. 2014;3(5):610–619.

29. Velcheti V, Schrump D, Saunthararajah Y. Ultimate Precision: Targeting Cancer but Not Normal Self-replication. Am Soc Clin Oncol Educ Book. 2018(38):950–963.

30. Habuka M, Fagerberg L, Hallstrom BM, Ponten F, Yamamoto T, Uhlen M. The Urinary Bladder Transcriptome and Proteome Defined by Transcriptomics and Antibody-Based Profiling. PLoS One. 2015;10(12):e0145301.

31. Yu Z, Liao J, Chen Y, et al. Single-Cell Transcriptomic Map of the Human and Mouse Bladders. J Am Soc Nephrol. 2019;30(11):2159–2176.

32. Kim J, Woo AJ, Chu J, et al. A Myc network accounts for similarities between embryonic stem and cancer cell transcription programs. Cell. 2010;143(2):313–324.

33. Lerdrup M, Johansen JV, Agrawal-Singh S, Hansen K. An interactive environment for agile analysis and visualization of ChIP-sequencing data. Nat Struct Mol Biol. 2016;23(4):349–357.

34. Cancer Genome Atlas Research N. Comprehensive molecular characterization of urothelial bladder carcinoma. Nature. 2014;507(7492):315–322.

35. Nickerson ML, Witte N, Im KM, et al. Molecular analysis of urothelial cancer cell lines for modeling tumor biology and drug response. Oncogene. 2017;36(1):35–46.

36. Kishtagari A, Ng KP, Jarman C, et al. A First-in-Class Inhibitor of ISWI-Mediated (ATP-Dependent) Transcription Repression Releases Terminal-Differentiation in AML Cells While Sparing Normal Hematopoiesis. Blood. 2018;132.

37. Oyama Y, Shigeta S, Tokunaga H, et al. CHD4 regulates platinum sensitivity through MDR1 expression in ovarian cancer: A potential role of CHD4 inhibition as a combination therapy with platinum agents. PLoS One. 2021;16(6):e0251079.

38. Jevtic Z, Matafora V, Casagrande F, et al. SMARCA5 interacts with NUP98-NSD1 oncofusion protein and sustains hematopoietic cells transformation. J Exp Clin Cancer Res. 2022;41(1):34.

39. Zikmund T, Paszekova H, Kokavec J, et al. Loss of ISWI ATPase SMARCA5 (SNF2H) in Acute Myeloid Leukemia Cells Inhibits Proliferation and Chromatid Cohesion. Int J Mol Sci. 2020;21(6).

40. Chen C, Dorado Garcia H, Scheer M, Henssen AG. Current and Future Treatment Strategies for Rhabdomyosarcoma. Front Oncol. 2019;9:1458.

41. Plagens RN, Tirado CSR, Li S, et al. Mapping the FOXA1 Interactome in ER+ Breast Cancer Cells Using Proximity Labeling Reveals Novel Interactions with the Orphan Nuclear Receptor NR2C2. Mol Cancer Res. 2025;23(12):997–1011.

42. Guo C, Zhang Y, Tan R, et al. Arid1a regulates bladder urothelium formation and maintenance. Dev Biol. 2022;485:61–69.

43. Agaimy A, Bertz S, Cheng L, et al. Loss of expression of the SWI/SNF complex is a frequent event in undifferentiated/dedifferentiated urothelial carcinoma of the urinary tract. Virchows Arch. 2016;469(3):321–330.

44. Tang HM, Kuay KT, Koh PF, et al. An epithelial marker promoter induction screen identifies histone deacetylase inhibitors to restore epithelial differentiation and abolishes anchorage independence growth in cancers. Cell Death Discov. 2016;2:16041.

45. Giannopoulou AF, Velentzas AD, Konstantakou EG, et al. Revisiting Histone Deacetylases in Human Tumorigenesis: The Paradigm of Urothelial Bladder Cancer. Int J Mol Sci. 2019;20(6).

46. von Knebel Doeberitz N, Paech D, Sturm D, Pusch S, Turcan S, Saunthararajah Y. Changing paradigms in oncology: Toward noncytotoxic treatments for advanced gliomas. Int J Cancer. 2022;151(9):1431–1446.

47. Gu X, Hu Z, Ebrahem Q, et al. Runx1 regulation of Pu.1 corepressor/coactivator exchange identifies specific molecular targets for leukemia differentiation therapy. J Biol Chem. 2014;289(21):14881–14895.

48. Gu X, Ebrahem Q, Mahfouz RZ, et al. Leukemogenic nucleophosmin mutation disrupts the transcription factor hub that regulates granulomonocytic fates. J Clin Invest. 2018;128(10):4260–4279.

49. Zavras PD, Shastri A, Goldfinger M, Verma AK, Saunthararajah Y. Clinical Trials Assessing Hypomethylating Agents Combined with Other Therapies: Causes for Failure and Potential Solutions. Clin Cancer Res. 2021;27(24):6653–6661.

