## Supplemental Data for "Urothelial-lineage master transcription factor hub proteomics shows mechanisms impeding urothelial cancer cell differentiation"

### SUPPLEMENTARY MATERIAL

**Table S1 (related to Figure 1) (Excel file). IP-LCMS/MS of IgG isotype control, FOXA1, CEBPB total spectral counts**

**Table S2 (related to Figure 1) (Excel file). Total spectral count comparison IP FOXA1 and CEBPB versus IP IgG isotype control**

**Table S3 (related to Figure 2). Umbrella cell and replication gene sets**

**Table S4 (related to Figure 2). Expression of umbrella and replication genes in primary UCs (TCGA RNA-seq data analyzed)**

**Table S5 (related to Figure 2). Gene expression by RNA-seq of Empty Vector vs ARID1A-transduced UC-6 cells (normalized counts)**

**Table S6 (related to Figure 2). Differentially expressed genes ARID1A transduction versus Empty Vector transduction (results of DESeq2 analyses)**

**2 supplementary figures**

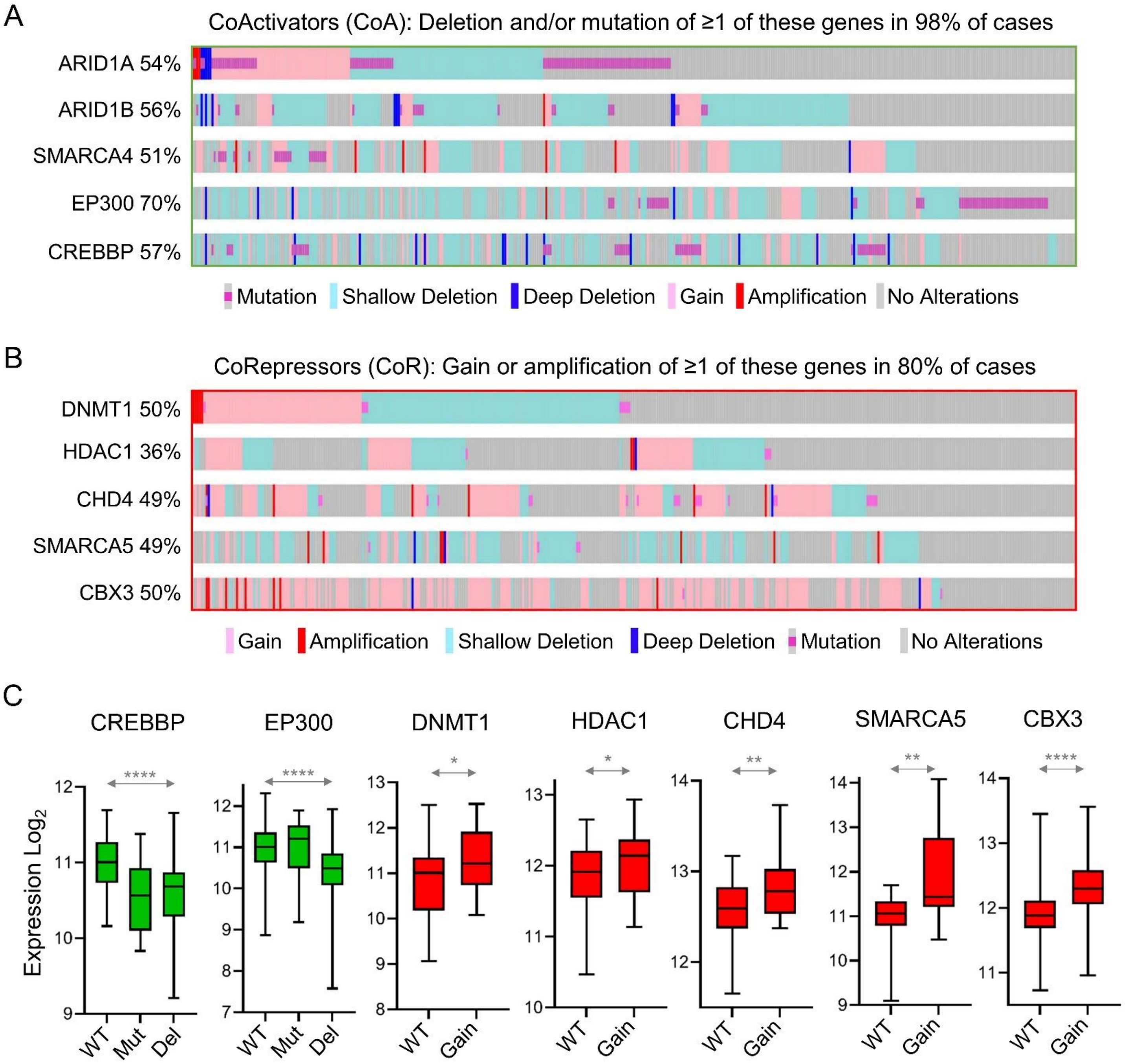


**Figure S1. A) CoAs that write or read (as part of the SWI/SNF complex) the H3K27ac activation mark deleted or mutated in 98% of UCs** (analyses of TCGA database[1]). These CoAs were prominent in the FOXA1/CEBPB interactomes in the IP-LCMS/MS analyses. Non-silent mutation and copy number calling as per TCGA[1], gene-level copy number estimates generated by the GISTIC2 method were thresholded to estimated values with -2 and -1 representing homozygous (deep) or single copy (shallow) deletion, 0 representing diploid normal copy, and 1 or 2 representing low-level (gain) or high-level copy number amplification. UCs n=397. **B) CoRs identified in the FOXA1/CEBPB interactomes were gained or amplified in 80% of UCs.** Analyses as per panel A. **C) Mutations and copy number deletions of CoA genes, and copy number gains and amplifications of CoR genes, in UCs decreased and increased expression of the genes respectively.** Copy number analyses as per panel A and B. Gene-level transcription estimates by RNA-Sequencing were analyzed as $\log_2(x+1)$ transformed RSEM normalized counts. Median+/-interquartile range, Mann-Whitney test 2-sided, *$p<0.05$, **$p<0.01$, ***$p<0.001$, ****$p<0.0001$. UCs n=397.

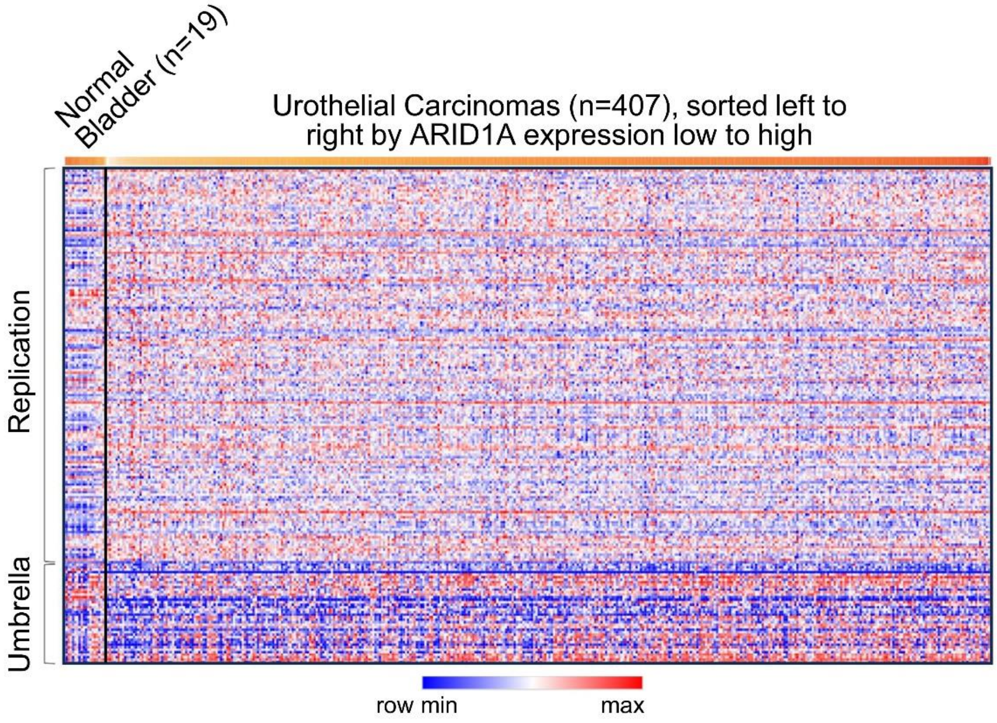


**Figure S2. Umbrella gene expression was lower, and replication gene expression higher, in primary UC tissue compared to non-cancerous bladder tissue** (TCGA RNA-sequencing, Table S4, primary UC samples n=407, non-cancer bladder n=19.

1. Cancer Genome Atlas Research N. Comprehensive molecular characterization of urothelial bladder carcinoma. *Nature*. 2014;507(7492):315-322.
